# Free Energy Differences from Molecular Simulations: Exact Confidence Intervals from Transition Counts

**DOI:** 10.1101/2022.11.19.517189

**Authors:** Pavel Kříž, Jan Beránek, Vojtěch Spiwok

## Abstract

Here we demonstrate a method to estimate the errors of free energy differences calculated by molecular simulations. The widths of the confidence intervals can be calculated solely from temperature and the number of transitions between states. Accuracy better than ± 4.184 kJ/mol (1 kcal/mol) can be achieved by a simulation at 300 K with four forward and four reverse transitions. Markovianity of the process is a pre-requisite. For a two-state Markovian system, the confidence interval suggested below is exact (not only asymptotic or approximative), regardless the number of transitions.

**TOC Graphic:** 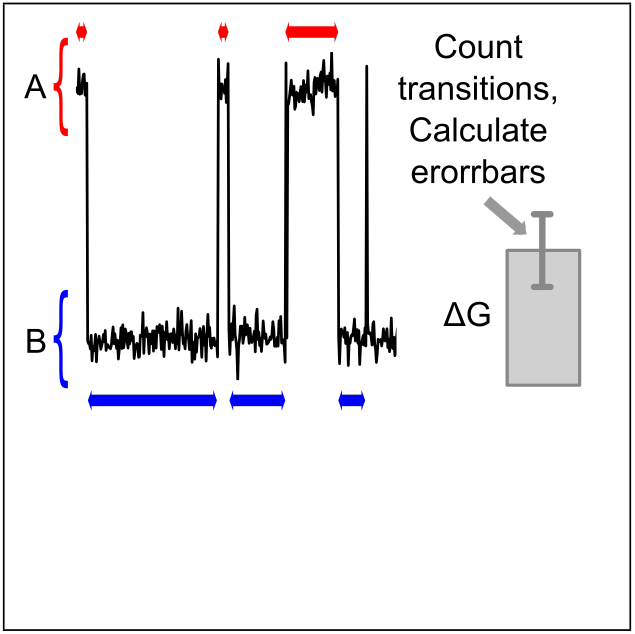

---

One of the main purposes of molecular simulations is to predict equilibrium constants and associated free energy differences for processes such as conformational changes, formation of non-covalent complexes, chemical reactions, or phase transitions. The equilibrium constant of a transition from state A to B can be determined experimentally as a fraction of the concentrations of B and A in equilibrium. In molecular simulations, it is common to simulate only one copy of the system (e.g., a single solvated protein or a protein-ligand pair), thus the equilibrium constant can be predicted as a fraction of time spent in states B and A. For practical application of such predictions, it is necessary to assess their accuracy. Current statistical methods used in this field are based on the treatment of autocorrelation.^1–3^ As an alternative, we present a simple method “JumpCount” based solely on the number of observed transitions.

First, we will derive an expression of the errors of an equilibrium constant and the associated free energy difference for a simple reversible transition from state A to state B. A simulation can be performed in a way that undergoes the same number of transitions from A to B (*n_A_*) and from B to A (*n_B_*). This is illustrated in Figure 1 for the number of transitions *n_A_* = *n_B_* = 3.

**Figure 1:**
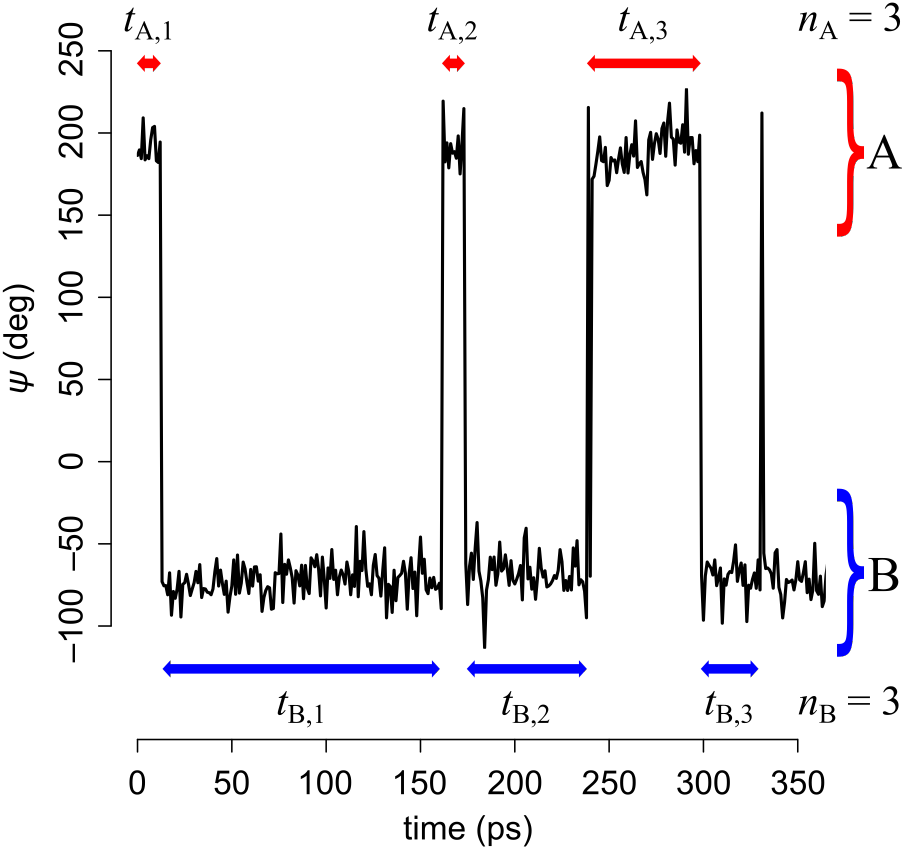
A schematic representation of a trajectory with three A to B and three B to A transitions (data taken from glycerol simulation).

For a Markovian process, the first time passage times for the transition from A to B (and vice versa) are exponentially distributed as *P*(*t_A_*) = *k*_1_ exp(-*k*_1_*t_A_*), where *k*_1_ is a rate constant for the transition from A to B. Analogously, *P*(*t_B_*) = *k*_−1_ exp(-*k*_-1_*t_B_*). It is possible to estimate *k*_1_ and *k*_-1_ as *n_A_*/∑_*i*_ *t_A,i_* and *n_B_*/∑_*i*_ *t_B,i_*, respectively.

The sum of independent random variables with exponential distributions follows the gamma distribution, namely Gamma(shape = *n_A_*, rate = *k*_1_) for ∑_*i*_ *t_A,i_*. The equilibrium constant 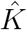 can be estimated as ∑_*i*_ *t_A,i_*/∑_*j*_ *t_B,j_* (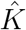 will be used as a symbol for an estimate of the true equilibrium constant *K*). The corresponding free energy difference can be estimated as 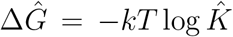. The fraction of the estimate and the true value of 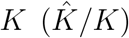 can be written as a rescaled fraction or independent gamma-distributed random variables and follows the Fisher-Snedecor F-distribution with degrees of freedom *d*_1_ = 2*n_B_* and *d*_2_ = 2*n_A_*. The confidence interval (CI) of *K* can therefore be calculated using the quantile function of the Fisher-Snedecor F-distribution. The 95-% of *K* CI can be calculated as:

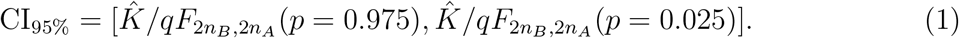

As a result, the CI of the free energy difference depends solely on temperature and the number of transitions. For 300 K, the CI for the free energy difference is 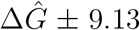 kJ/mol (2.18 kcal/mol) for *n_A_* = *n_B_* = 1, 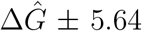 kJ/mol (1.35 kcal/mol) for *n_A_* = *n_B_* = 2, etc. (see Table 1). The confidence interval 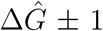 kcal/mol, which is often used as a threshold of accuracy in molecular simulations, can be reached in a simulation with *n* = 4.

**Table 1:** 95 % confidence intervals at 300 *K* (mean ± error) for processes with numbers of transitions from A to B (*n_A_*) and from B to A (*n_B_*).

| $n_A$ | $n_B$ | error<br>(kJ/mol) | error<br>(kcal/mol) | $n_A$ | $n_B$ | error+<br>(kJ/mol) | error+<br>(kcal/mol) | error-<br>(kJ/mol) | error-<br>(kcal/mol) |
| --- | --- | --- | --- | --- | --- | --- | --- | --- | --- |
| 1 | 1 | 9.14 | 2.18 | 2 | 1 | 5.90 | 1.41 | 9.15 | 2.19 |
| 2 | 2 | 5.64 | 1.35 | 3 | 2 | 4.56 | 1.09 | 5.53 | 1.32 |
| 3 | 3 | 4.39 | 1.05 | 4 | 3 | 3.83 | 0.92 | 4.30 | 1.03 |
| 4 | 4 | 3.71 | 0.89 | 5 | 4 | 3.37 | 0.80 | 3.64 | 0.87 |
| 5 | 5 | 3.27 | 0.78 | 6 | 5 | 3.03 | 0.72 | 3.21 | 0.77 |
| 6 | 6 | 2.96 | 0.71 | 7 | 6 | 2.78 | 0.66 | 2.91 | 0.69 |
| 7 | 7 | 2.72 | 0.65 | 8 | 7 | 2.58 | 0.62 | 2.68 | 0.64 |
| 8 | 8 | 2.53 | 0.61 | 9 | 8 | 2.42 | 0.58 | 2.49 | 0.60 |
| 9 | 9 | 2.38 | 0.57 | 10 | 9 | 2.29 | 0.55 | 2.34 | 0.56 |
| 10 | 10 | 2.25 | 0.54 | 11 | 10 | 2.17 | 0.52 | 2.22 | 0.53 |
| 20 | 20 | 1.57 | 0.37 | 21 | 20 | 1.54 | 0.37 | 1.55 | 0.37 |

The concept described above can be easily generalized to *n_A_* = *n_B_* + 1 (we denote the starting state as A). In this case, the confidence interval can be calculated by setting *df*_1_ = *n_B_* and *df*_2_ = *n_A_* in the F-distribution. The confidence interval is then asymmetric (see Table 1).

It is also possible to generalize the concept to binding, such as simulation of proteinligand interactions. The dissociation constant of a complex PL (the equilibrium constant of PL ⇌ P + L) can be expressed as *K_d_* = ∑(*c_L_*∑*t_P_*)/*t_PL_*, where *c_L_* is the concentration of the ligand in the simulation box in the unbound state. The accuracy of ∑*t_P_*/∑*t_PL_*, (and thus for the resulting *K_d_* and binding Δ*G_bind_*) can be calculated as described above.

The easiest way to test the concept is to generate the first time passage times as exponentially distributed random numbers. This was performed in the R software (see Supporting Information). We tested scenarios with *n_A_* and *n_B_* set to 1 to 20 and with *K* set to 1 to 1,000. The values of *k*_-1_ and *k*_1_ were set to 1 and *K*, respectively (in arbitrary units). For each pair of *n* and *K* we generated 10,000 sets of first time passage times and calculated 95-% CI of *K* and 95-% CI of the free energy difference. We expect the rate of type I errors (i.e., the fraction of trials for which the predefined *K* lies outside the calculated CI) to be 5 %. Indeed, the type I error rate ranged from 4.64 to 5.60 % with a median of 5.03 %. Similar results were observed for nA = *n_B_* + 1 (see Supporting Information).

For a system with multiple states (e.g., A, B, and C), it is necessary to count only the accomplished transitions between states. For example, when calculating Δ*G_A→C_* it is necessary to count the process with transitions A → B → A → B → Casa single accomplished transition from A to C. The resulting numbers of accomplished transitions *n_A_* and *n_C_* can be used in Equation 1 to obtain CI of *K_A→C_* and Δ*G_A→C_* (see Supporting Information for a demonstration of the fact that the sum of *t_Ai_* and the sum of *t_Ci_* follow the Gamma(shape = *n_A_*,…) and Gamma(shape = *n_C_*,…) distributions, respectively).

As an alternative, for a system with multiple states, it is possible to calculate the free energy differences for states with direct transitions (e.g., A to B and B to C, for a system with A ⇌ B ⇌ C transitions) separately and combine errors as (∑*error s*^2^)^1/2^. This approach is used for variables with normal distribution. It is possible to apply it to systems with a high number of transitions, which guarantees a close-to-normal distribution of 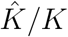.

An example of a molecular system with multiple states is a glycerol molecule in water.^4^ Each of the two O-C-C-O torsion angles can adopt three minima. This gives nine combinations; however, the three pairs are equivalent as a result of the symmetry of the molecule; thus, six conformers can be experimentally resolved. Equilibria of these conformers have been studied by molecular simulations as well as experimentally.^4,5^

Here, we performed a 1 μs simulation (see Supporting Information for details) of glycerol in water and calculated the equilibria for six conformers. Consistent with simulation and experimental studies^4,5^ we observed conformer populations *αγ* > *αβ* > *αα* > *βγ* > *γγ* > *ββ* ~ *γγ*. Confidence intervals for all conformers relative to *αγ* were calculated at times 5, 10, 20, 50, 100, 200, 500, and 1,000 ns. In total, 37 confidence intervals were calculated (eight for each conformer except *αγ*, confidence intervals for *γγ* were not available at 5, 10, and 20 ns due to no sampling). These confidence intervals were compared with the value of Δ*G* calculated from the whole trajectory. Since we do not know the exact value of Δ*G*, we used this value as a “ground truth”.

One of these 37 intervals was not spanning the ground truth Δ*G*. This was the case of Δ*G* of *βγ* at 50 ns (Figure 2). The distance of Δ*G* from the confidence interval was very low. Figures for other conformers are available in Supporting Information. Since we compare the confidence intervals with the estimate of Δ*G* calculated for whole trajectory, not the exact value of Δ*G*, we expect the rate of type 1 errors to be lower than 5 %. This is in agreement with the observed 1/37 = 2.7 %.

**Figure 2:**
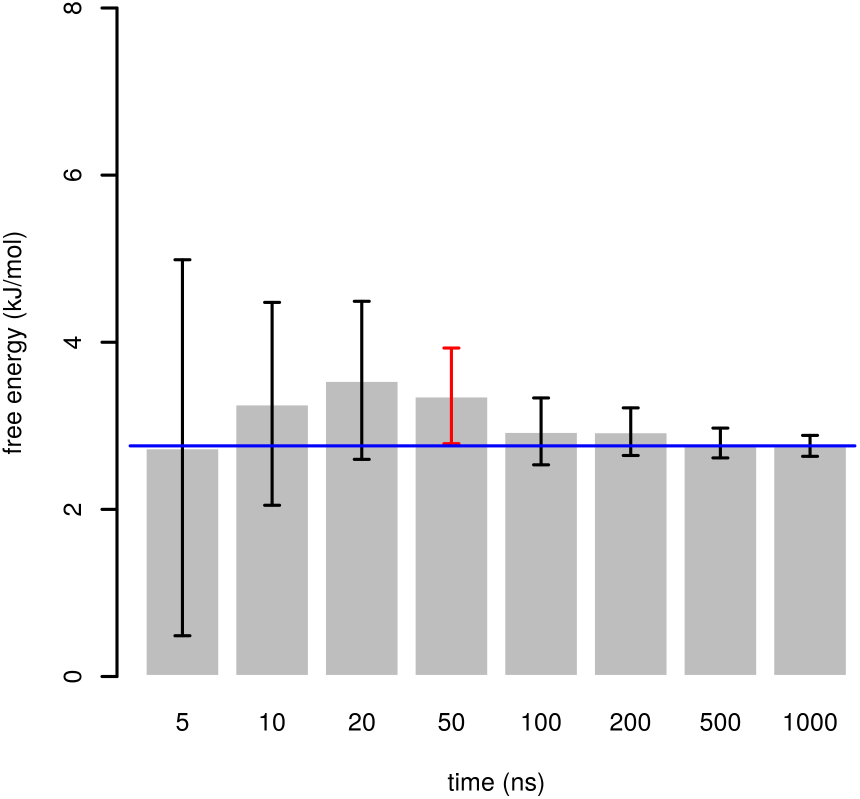
Confidence intervals of free energy of *βγ* conformer of glycerol, relative to *αγ*. The value of free energy calculated for the whole simulation is depicted as a blue line. A confidence interval that does not span this value is depicted in red.

The main prerequisite of the above-outlined approach is Markovianity of the processes studied. This is usually fulfilled for transitions associated with a single energy barrier, such as simple conformational changes or chemical reactions. More complex transitions, such as protein folding, can be non-Markovian.^6^ The method of molecular dynamics simulation is Markovian because each state of the molecular system depends solely on the previous state, not on the history. However, coarse graining of the system representation into a few substates (e.g., folded and unfolded, ligand-bound and unbound) and ignoring the complex kinetics within these states may cause the Markovianity condition to be not fulfilled.

Keeping in mind the limitation of the Markovianity prerequisite, we applied our approach to the folding and unfolding trajectories of fast folding miniproteins.^7^ Trajectories were kindly provided by D.E. Shaw research. Root mean square deviation (RMSD) profiles from the native structure were calculated, and folded and unfolded states were assigned by visual inspection of RMSD profiles and trajectories. Folding free energies were estimated as 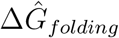 for the whole trajectories (multiple trajectories of the same system were combined). These values were used as a ground truth.

They were compared with 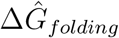 calculated at 20 points equally distributed along each trajectory. The ground truth was outside the 95 % confidence intervals for six of 432 (1.39 %) values. These confidence intervals are shown in Figure 3. Corresponding plots for other systems can be obtained in Supporting Information.

**Figure 3:**
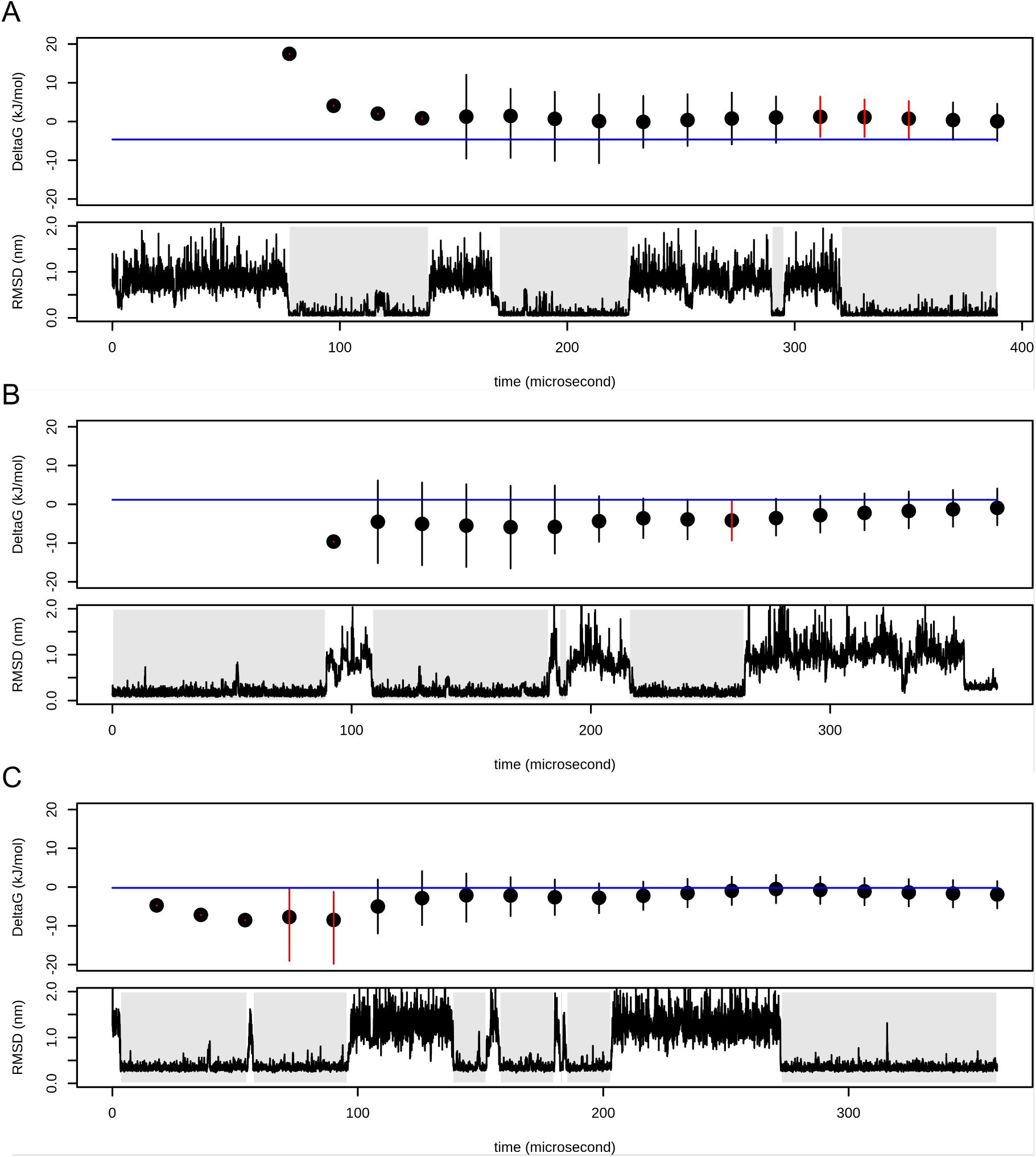
Confidence intervals of folding free energies for (A) NTL9 (simulation 2), (B) protein G (simulation 0), and (C) *α*3D (simulation 1). Top part of each subplot shows calculated folding free energy with confidence intervals depicted as error bars. The value of folding free energy calculated for the whole simulation is depicted as a blue line. Confidence intervals that do not span this value are depicted in red. Bottom parts show RMSD profiles. Folded states are highlighted by gray background.

Since we do not know the exact value of Δ*G_folding_* and we used the value calculated for the whole simulation as the ground truth, we expect a lower number of type 1 errors (values outside 95 % CI) than 5 %, which is in good agreement with the observed 1.39 %. Furthermore, all values outside the confidence intervals were located very close to them.

The results show that our approach can be applied to fast-folding miniproteins. Either the degree of non-Markovianity in these systems is not high enough to significantly affect the performance of our approach or this approach is robust enough for the non-Markovianity typical for biomolecular systems.

The problem of non-Makovianity can be solved by dissection of sampled states into a minimal set of substates for which mutual transitions are Markovian. This approach is used when building Markov state models.^8^ Alternatively, it would be possible to estimate the true distribution of *t_A_* and tB for non-Markovian systems and derive corresponding ditributions for *K* and Δ*G*.

We argue that the approach can be generalized to biased simulations with a static bias potential, such as umbrella sampling.^9^ Adapting the approach to methods with a time-dependent bias potential, such as metadynamics,^10^ will be the subject of future research.

In Supporting Information, it is possible to find commands to calculate confidence intervals in various programming languages. Data necessary to reproduce all calculations are available online via Zenodo (DOI: 10.5281/zenodo.7337600). Online calculator of CI is available at https://jumpcount.cz.

## Acknowledgement

The authors would like to thank D.E. Shaw Research for providing us trajectories. The work was supported by the Czech Science Foundation (22-29667S and 19-16857S). Computational resources were supplied by the project “e-Infrastruktura CZ” (e-INFRA CZ LM2018140) and ELIXIR-CZ project (LM2018131) supported by the Ministry of Education, Youth and Sports of the Czech Republic.

## Supporting Information Available

Testing of the method on random numbers with exponential distribution, computational details and detailed results of simulation of aqueous glycerol, full results of application of the method on fast-folding mini-proteins and instructions to calculate errors in various programming languages and available as Supporting Information.

