## Supporting Information for "Free Energy Differences from Molecular Simulations: Exact Confidence Intervals from Transition Counts"

### Simulation using generated random numbers

Simulations using generated random numbers were performed in R. First, we wrote a function to generate a set of $t_A$ and $t_B$ using function `rexp`. It generated $n_A = n_B$ number of $t_A$ and $t_B$ with predefined values of $k_1$ set to $K$ and $k_{-1}$ set to 1. Next, CIs were calculated and returned to the output of the function.

The function was applied 10,000 times for each combination of $n_A = n_B$ from 1 to 20 and $K$ set to 1, 2, 5, 10, 20, 50, 100, 200, 500 and 1000. Fractions of trials for which $K$ is located outside CI are plotted in the Figure S1 as a heatmap. The results are in good agreement with the expected rate of type 1 errors (5 % for 95-% CI).

Figure S1: Rates of type 1 errors (in %) for different $n_A = n_B$ and $K$ in simulations using generated random numbers with exponential distribution.

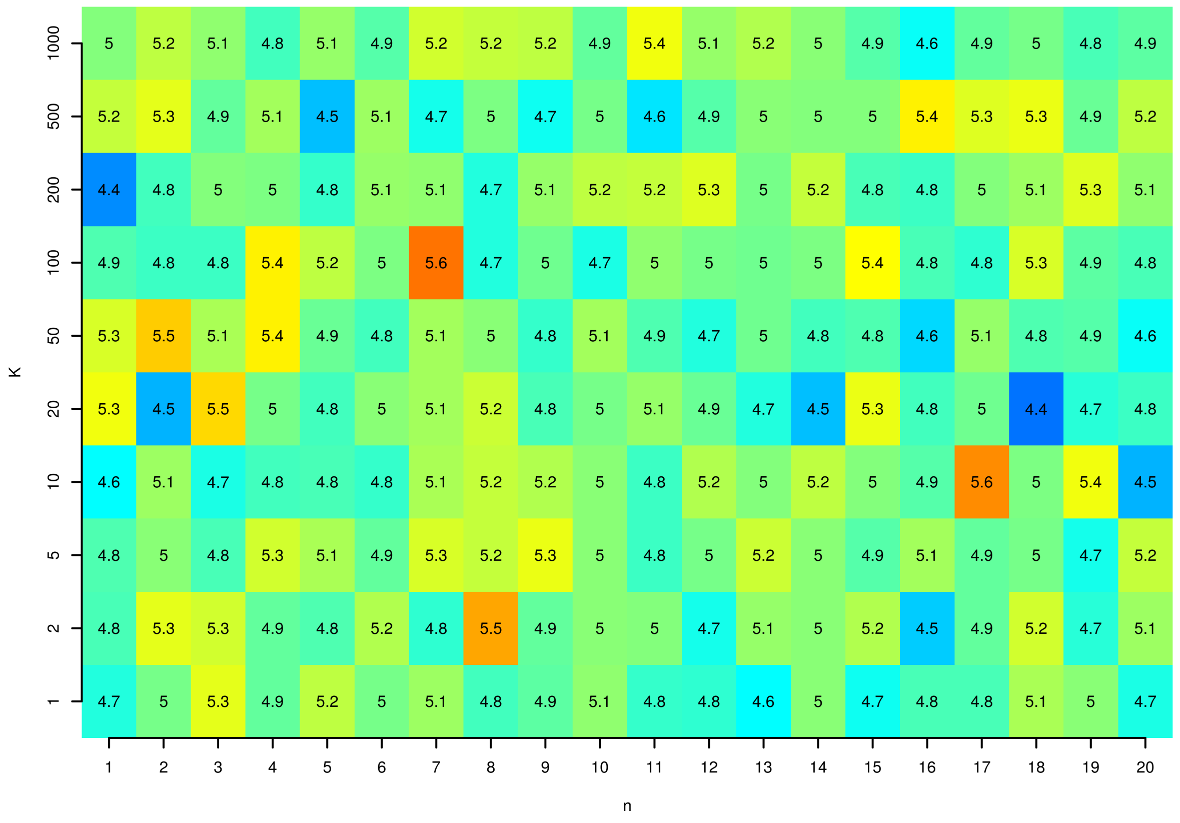

The same simulations as described above were performed with $n_A = n_B + 1$. The function was applied 10,000 times for each combination of $n$ ($n_A = n_B + 1 = n + 1$) from 1 to 20 and $K$ set to 1, 2, 5, 10, 20, 50, 100, 200, 500 and 1000. Fractions of trials for which $K$ is located outside CI are plotted in the Figure S2 as a heatmap. Again, the results are in good agreement with the expected rate of type 1 errors (5 % for 95-% CI).

Figure S2: Rates of type 1 errors (in %) for different $n$ ($n_A = n_B + 1 = n + 1$) and $K$ in simulations using generated random numbers with exponential distribution.

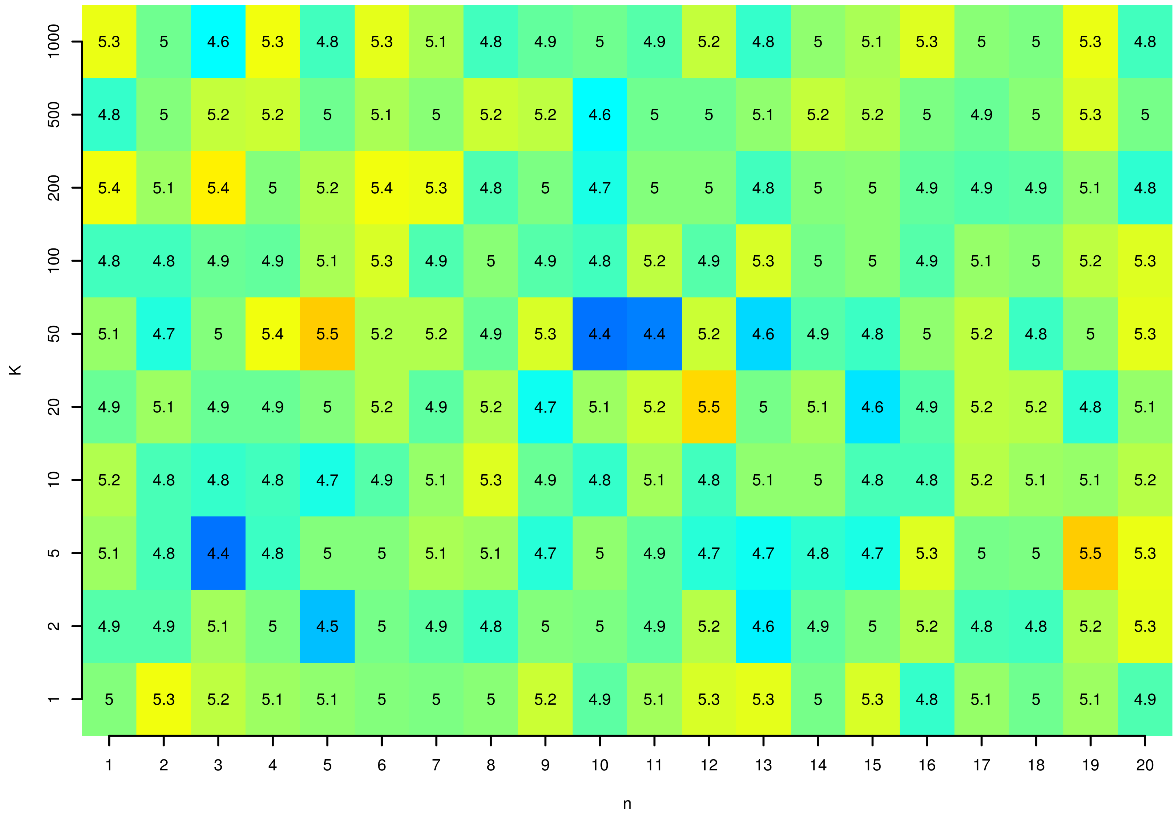

The reasoning that Equation 1 is correct also for the three-state system with transitions A $\rightleftharpoons$ B $\rightleftharpoons$ C (with $n_A$ and $n_C$ being the numbers of accomplished transitions from A to C and from C to A, respectively): Take an example of a continuous-time Markov chain with possible transitions A $\to$ B with the rate constant $k_1 = K_{AB} > 0$, B $\to$ C with the rate constant $k_2 = K_{BC} > 0$, C $\to$ B with the rate constant 1 and B $\to$ A with the rate constant 1 (rate constants equal to equilibrium constants and 1 were used for simplicity; however, it can be generalized for any values).

Furthermore, consider an example of a single accomplished transition A $\to$ B $\to$ A $\to$ B $\to$ C, with the number of visits of state A being $N_A = 2$ and the total time spent in state A being $T_A = t_{A1} + t_{A2}$, where $t_{Ai}$ is the occupation time in state A during $i$-th visit.

Firstly, note that the number of visits $N_A$ of state A within a single acomplished transition is a random variable having geometric distribution Ge$[p = k_2/(1+k_2)]$. This is because every time the process visits state B, it moves either to state A (with probability $1/(1 + k_2)$) or to state C (with probability $k_2/(1 + k_2)$).

Next, individual occupation times $t_{Ai}$ are independent random variables that have an exponential distribution with the rate constant $k_1$, and they are independent of $N_A$. Now, the total occupation time in state A during a single accomplished transition $T_A = t_{A1} + t_{A2} + ... + t_{AN_A}$ has a compound distribution and we can show (using the machinery of moment-generating functions) that it is again an exponential distribution with intensity $k_1k_2/(1 + k_2)$. Similarly, we can demonstrate that the total occupation time in state C during a single accomplished transition from C to A has an exponential distribution with intensity $1/(1 + k_2)$.

Now the reasoning follows the same lines as in the 2-state case: sums of total occupation times in A and C over all accomplished transitions are independent and follow gamma distributions Gamma(shape $= n_A$, rate $= k_1k_2/(1 + k_2)$) and Gamma(shape $= n_C$, rate $= 1/(1 + k_2)$), respectively. Their ratio (denote $\hat{K}$) is the estimator of the stationary probabilities ratio, and the expression

$$\hat{K}\,\frac{n_A}{n_C}\,\frac{1}{k_1k_2} = \frac{\hat{K}}{K_{AB}K_{BC}} = \frac{\hat{K}}{K_{AC}}$$

has Fisher-Snedecor distribution $F(2n_C, 2n_A)$. From this, it easily follows that Equation (1) provides exact confidence interval for the unknown quantity $k_1k_2 = K_{AB}K_{BC} = K_{AC}$.

Simulations with three states with transitions A $\rightleftharpoons$ B $\rightleftharpoons$ C were also tested numerically. First, we wrote a function to generate a set of $t$. For each application of this function, the simulation started from the state A. The value of $t_{A\to B}$ was generated and the state changed to B. Next, values of $t_{B\to A}$ and $t_{B\to C}$ were generated. If $t_{B\to C} < t_{B\to A}$, the state changed to C, otherwise it changed to A. Analogously, a pair of $t$ was generated for any state until the desired number of accomplished transitions from A to C and C to A was observed. For example, it is necessary to count the process with transitions A $\to$ B $\to$ A $\to$ B $\to$ C as a single accomplished transition from A to C.

Confidence intervals for $K$ were calculated as:

$$\mathrm{CI}_{95\%} = [\hat{K}/qF_{2n_C,2n_A}(p = 0.975), \hat{K}/qF_{2n_C,2n_A}(p = 0.025)]. \qquad (1)$$

where $n_A$ and $n_C$ are the numbers of accomplished transitions from A to C and C to A, respectively.

The function was applied 10,000 times for each combination of $n_A = n_C$ set to 1, 2, 5, 10, 20 and 50 and $K_{A\to B}$ set to 10 and $K_{B\to C}$ was set to 1, 2, 5, 10 and 20.

Fractions of trials for which $K_{A\to C} = K_{A\to B}K_{B\to C}$ is located outside CI are plotted in the Figure S3 as a heatmap. Again, the results are in good agreement with the expected rate of type 1 errors (5 % for 95-% CI).

Figure S3: Rates of type 1 errors (in %) for different $n_A = n_B$ and $K_{A \to C}$ in simulations using generated random numbers with exponential distribution for a system with states A, B and C.

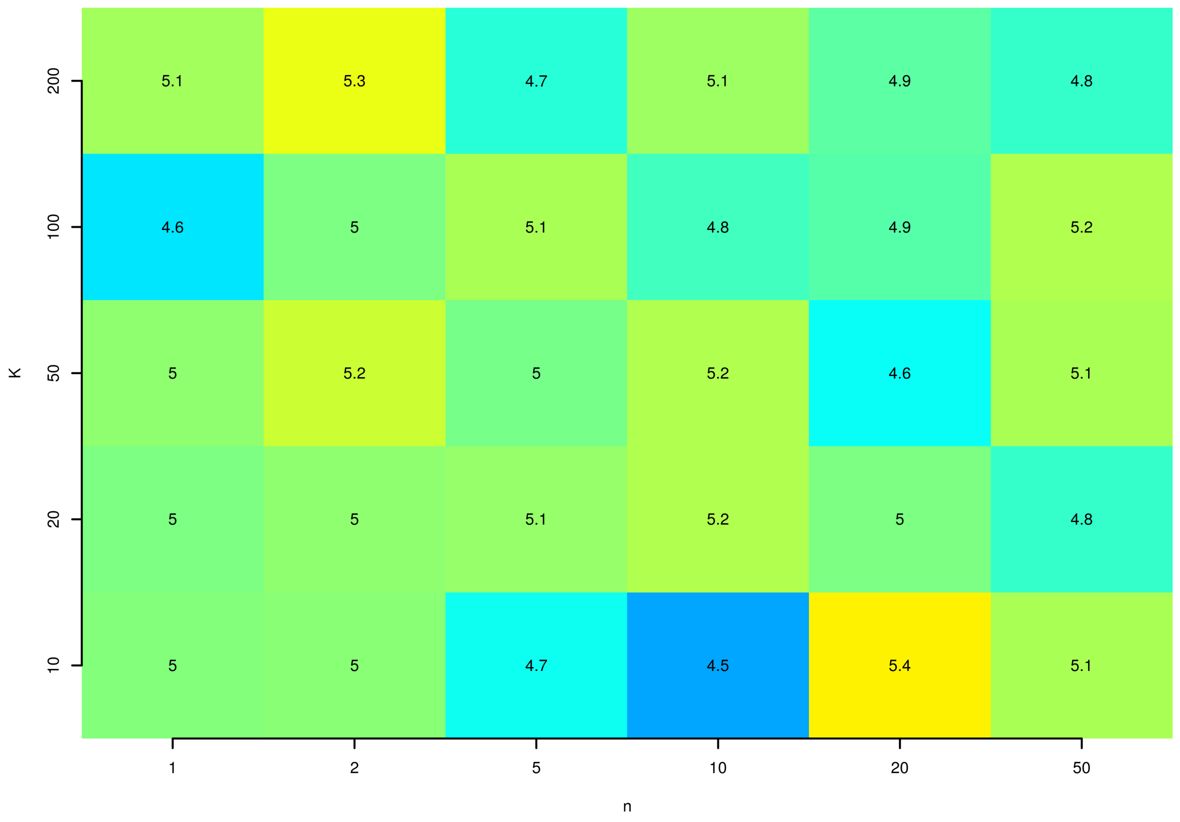

### Glycerol in Water

#### Computational details

MD simulation was conducted using GROMACS 2018.6. Relevant preparation steps were conducted using GROMACS 2022.3.[1]

The preparation of the system for MD simulation was as follows: Starting structure of *sn*-glycerol was obtained by conversion of SMILES file to PDB format using Open Babel.[2] Topology was built manually according to Glycam06.[3] Partial atomic charges were calculated at HF/6-31G*//HF/6-31G* level of the theory using the RESP method.[4] Simulation box was cubic with a size of 2.94013 nm. The molecule was solvated using `gmx solvate` with TIP3P water. No ions were added, as the system had zero net charge. Potential energy of the system was then minimized using the steepest descent integrator, until the maximum force acting on any atom was lower than 1000 kJ/mol/nm. This step was followed by isothermal-isochoric equilibration and isothermal-isobaric equilibration, each at 300 K for 100 ps. Then, 1 µs long MD simulation was conducted. At the beginning of the simulation, the velocities of atoms were generated randomly from Maxwell distribution for 300 K. In the relevant steps of equilibration and MD run, the following parameters were used: Leap-frog integrator (md), radius for short-range electrostatic and van der Waals was set to 1 nm (spanning the whole molecule of glycerol). Particle Mesh Ewald method[5] was used for computing long-range electrostatic interactions. Temperature coupling was conducted using Parrinello-Bussi thermostat[6] and pressure coupling was conducted with Parrinello-Rahman barostat.[7]

After obtaining the trajectory of glycerol from the MD run, the values of torsion angles in the molecule were computed at each 1 ps using `plumed driver`.[8]

#### Results

Figure S4: Torsion angles $\phi$ and $\psi$ sampled during the simulations.

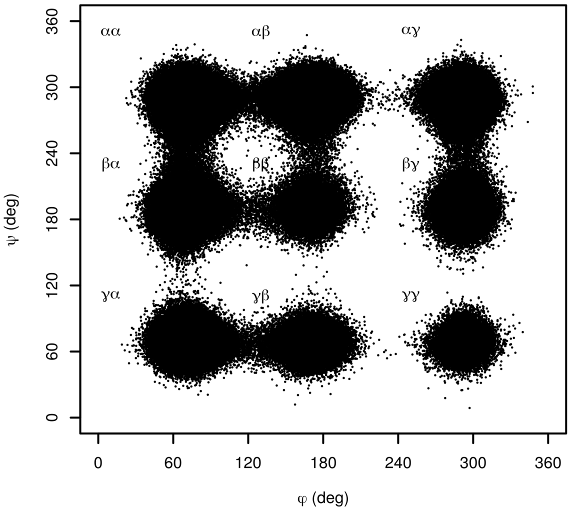

Figure S5: Confidence intervals of free energy of $\alpha\alpha$ conformer of glycerol, relative to $\alpha\gamma$. The value of free energy calculated for the whole simulation is depicted as a blue line.

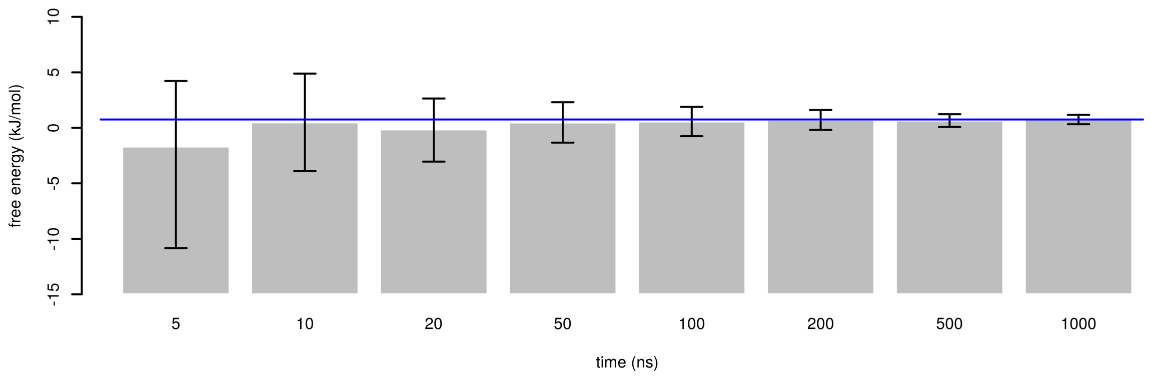


Figure S6: Confidence intervals of free energy of $\alpha\beta$ conformer of glycerol, relative to $\alpha\gamma$. The value of free energy calculated for the whole simulation is depicted as a blue line.

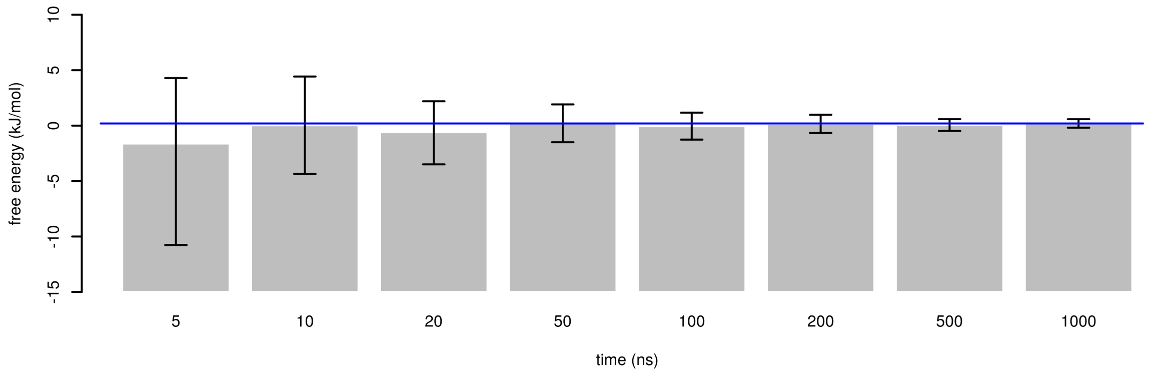

Figure S7: Confidence intervals of free energy of $\beta\beta$ conformer of glycerol, relative to $\alpha\gamma$. The value of free energy calculated for the whole simulation is depicted as a blue line.

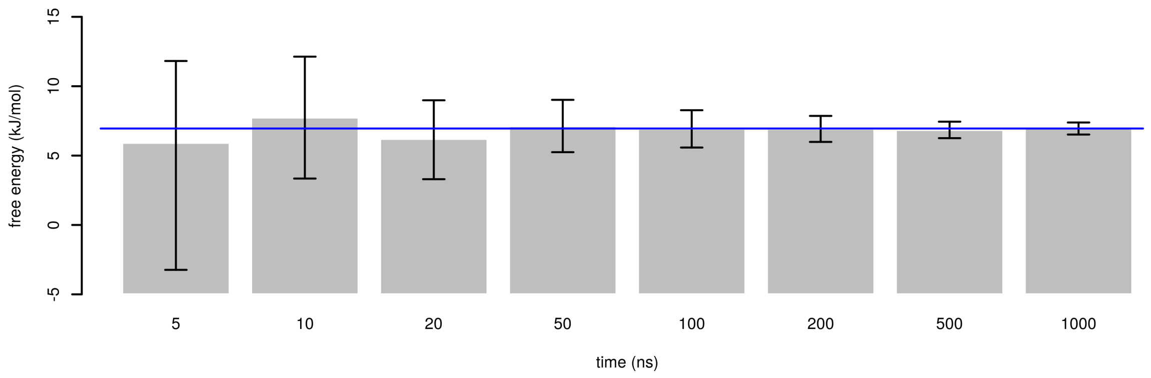


Figure S8: Confidence intervals of free energy of $\beta\gamma$ conformer of glycerol, relative to $\alpha\gamma$. The value of free energy calculated for the whole simulation is depicted as a blue line. A confidence interval that does not span this value is depicted in red.

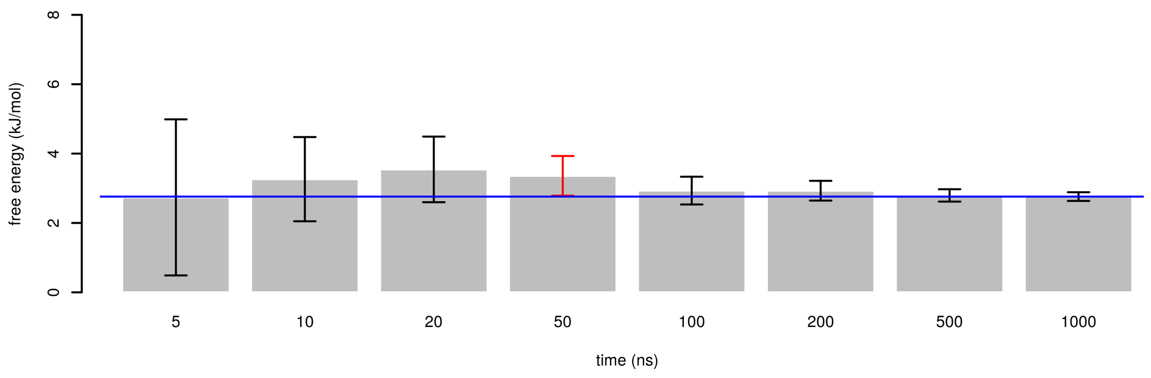

Figure S9: Confidence intervals of free energy of $\gamma\gamma$ conformer of glycerol, relative to $\alpha\gamma$. The value of free energy calculated for the whole simulation is depicted as a blue line.

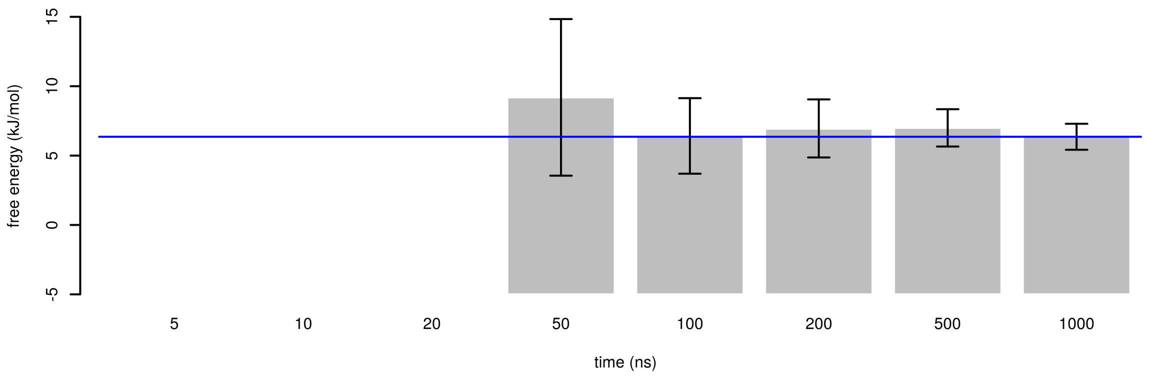

### Fast folding miniproteins

#### Results

Figure S10: Confidence intervals of folding free energy of Chignolin. The value of free energy calculated for the whole simulation is depicted as a blue line.

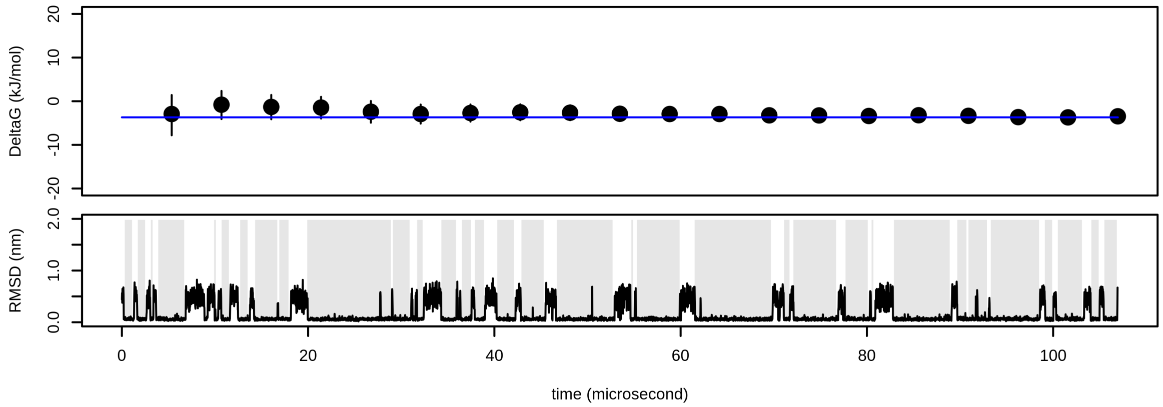


Figure S11: Confidence intervals of folding free energy of Trp-cage. The value of free energy calculated for the whole simulation is depicted as a blue line.

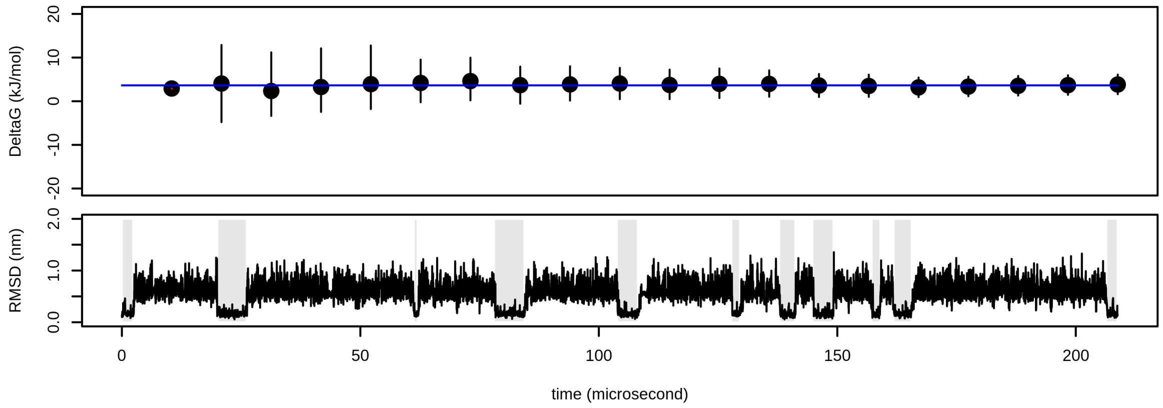

Figure S12: Confidence intervals of folding free energy of BBA (simulation 0). The value of free energy calculated from all simulations is depicted as a blue line.

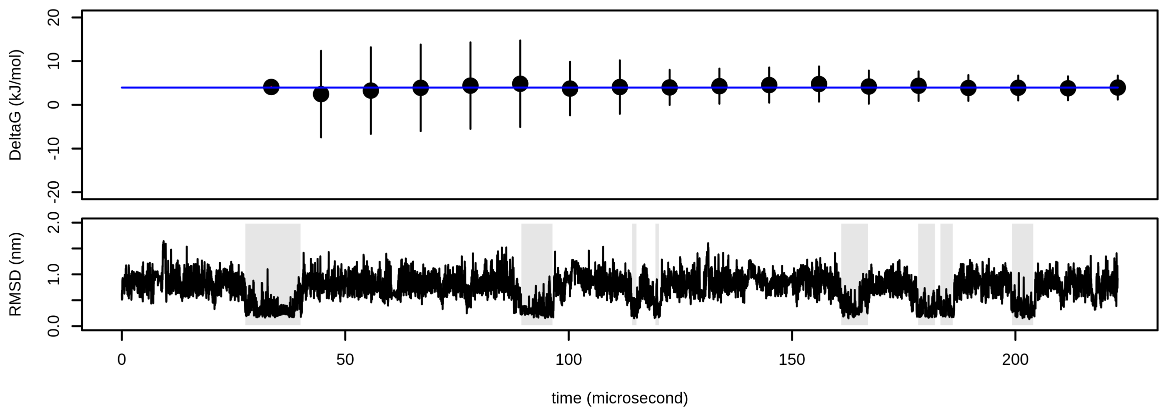


Figure S13: Confidence intervals of folding free energy of BBA (simulation 1). The value of free energy calculated from all simulations is depicted as a blue line.

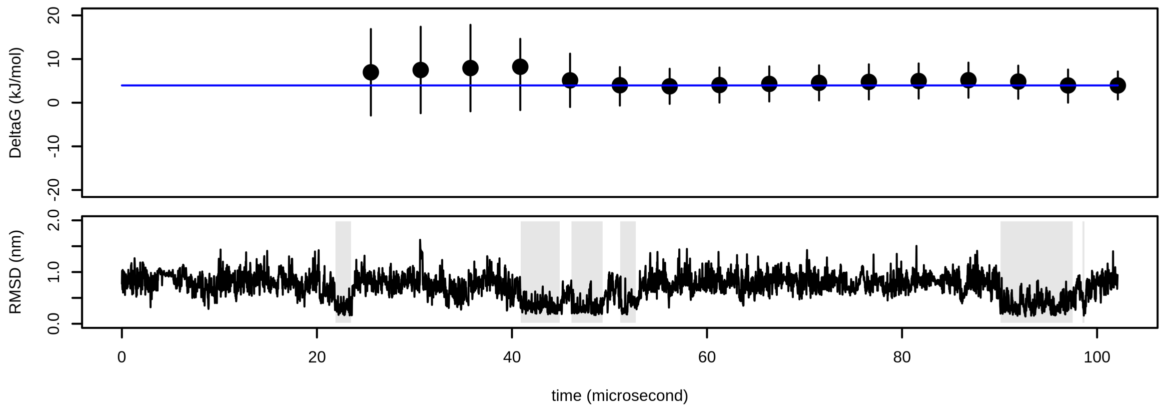

Figure S14: Confidence intervals of folding free energy of villin. The value of free energy calculated for the whole simulation is depicted as a blue line.

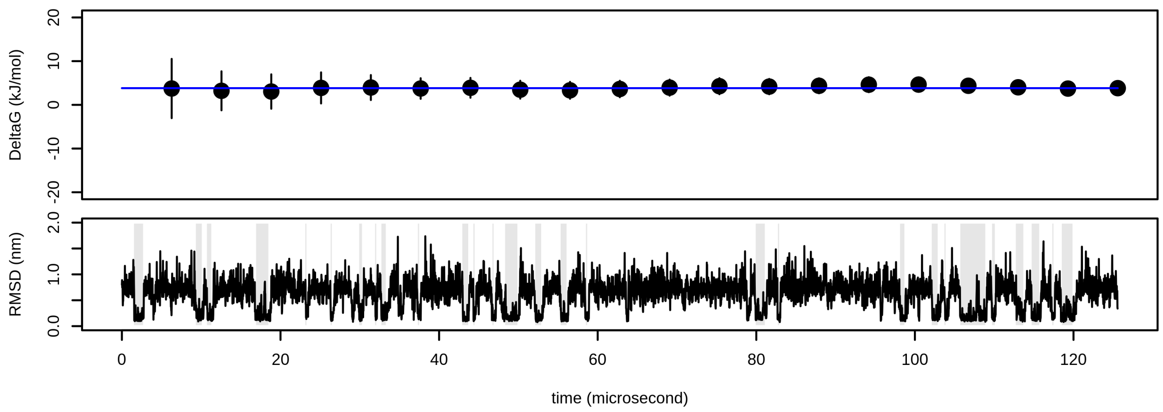


Figure S15: Confidence intervals of folding free energy of WW domain (simulation 0). The value of free energy calculated from all simulations is depicted as a blue line.

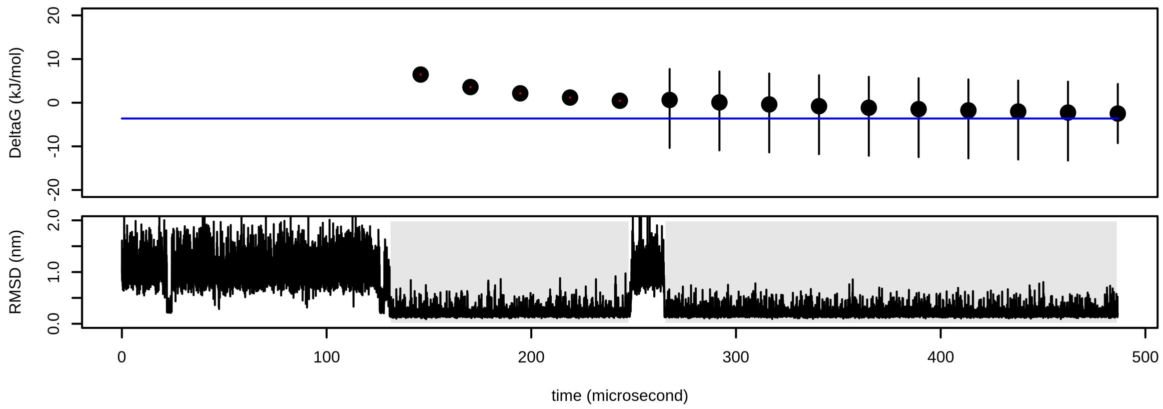

Figure S16: Confidence intervals of folding free energy of WW domain (simulation 1). The value of free energy calculated from all simulations is depicted as a blue line.

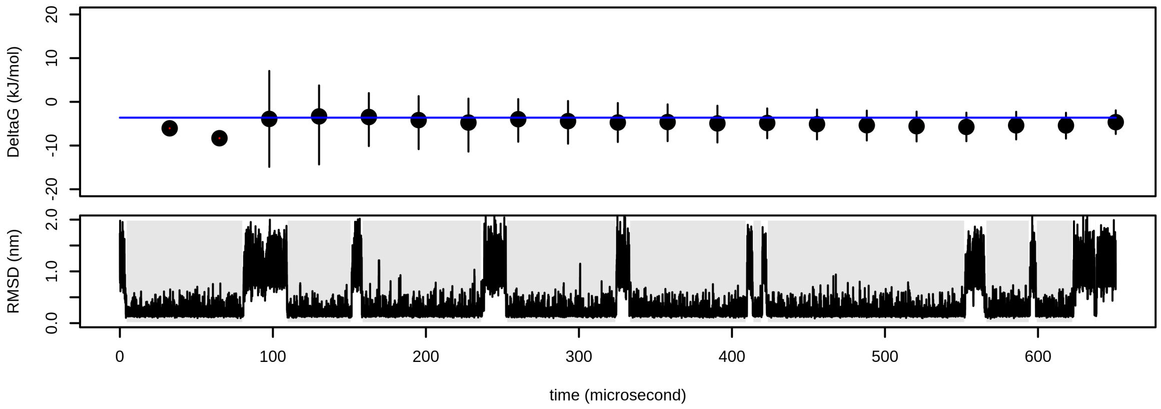


Figure S17: Confidence intervals of folding free energy of NTL9 (simulation 0). The value of free energy calculated from all simulations is depicted as a blue line.

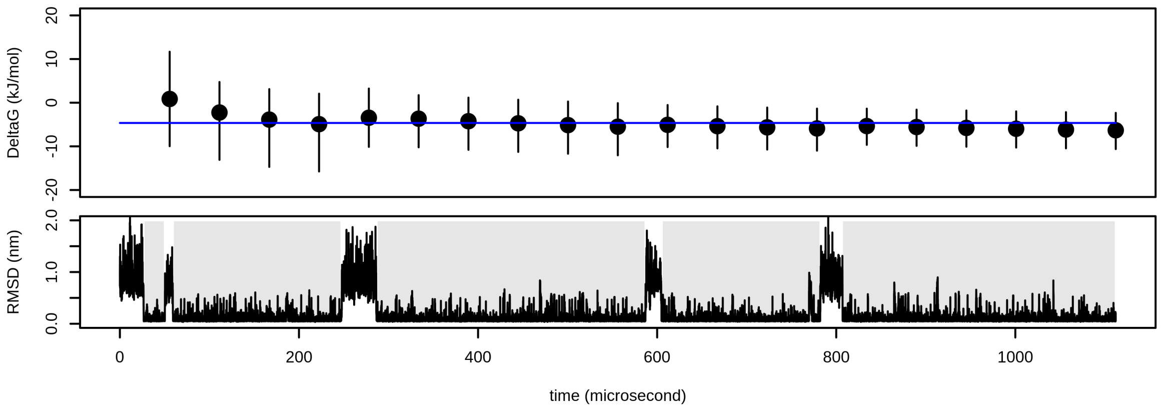

Figure S18: Confidence intervals of folding free energy of NTL9 (simulation 1). The value of free energy calculated from all simulations is depicted as a blue line.

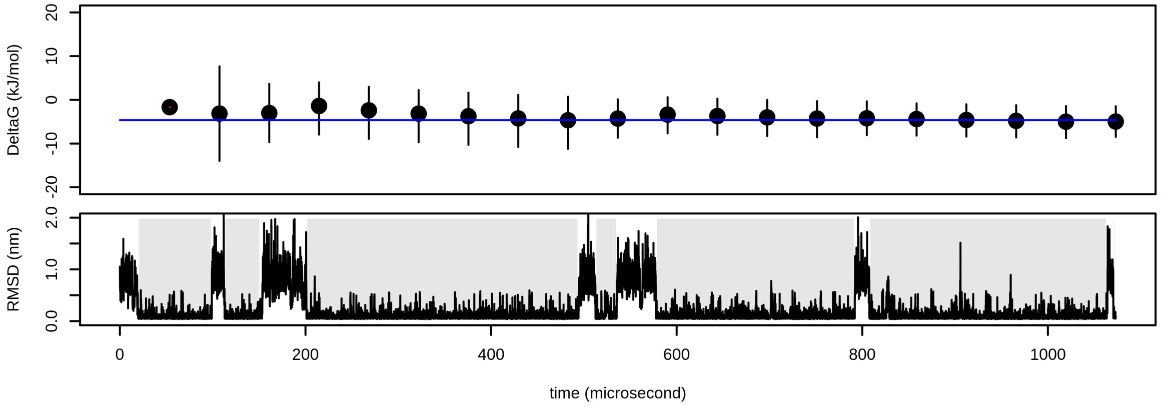


Figure S19: Confidence intervals of folding free energy of NTL9 (simulation 2). The value of free energy calculated from all simulations is depicted as a blue line. Confidence interval that do not span this value are depicted in red.

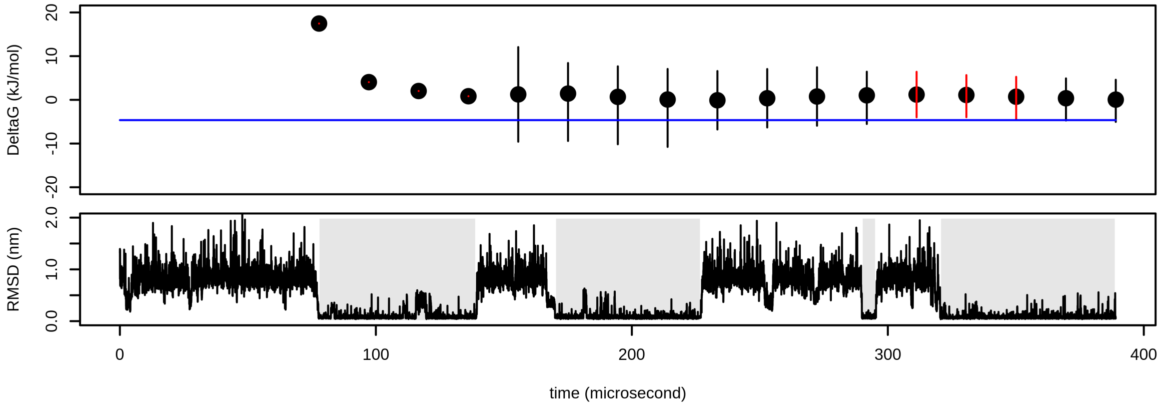

Figure S20: Confidence intervals of folding free energy of NTL9 (simulation 3). The value of free energy calculated from all simulations is depicted as a blue line.

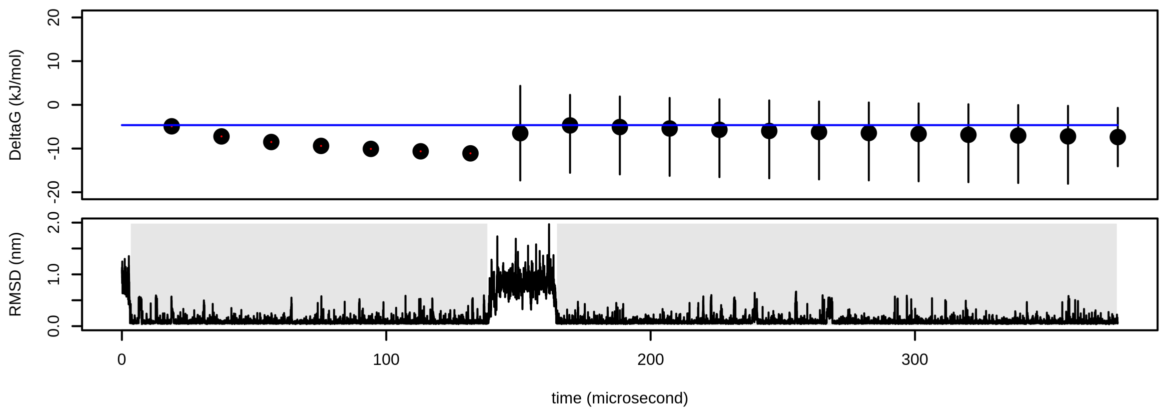


Figure S21: Confidence intervals of folding free energy of BBL (simulation 0). The value of free energy calculated from all simulations is depicted as a blue line.

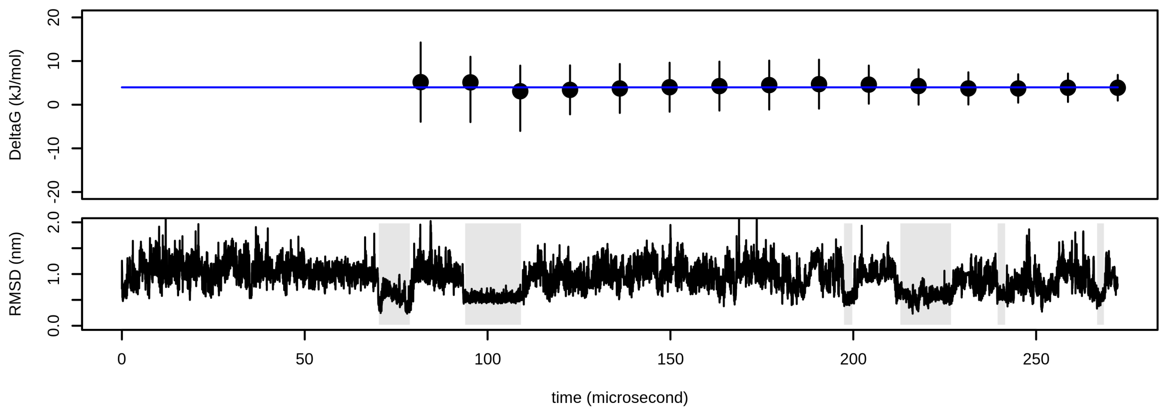

Figure S22: Confidence intervals of folding free energy of BBL (simulation 1). The value of free energy calculated from all simulations is depicted as a blue line.

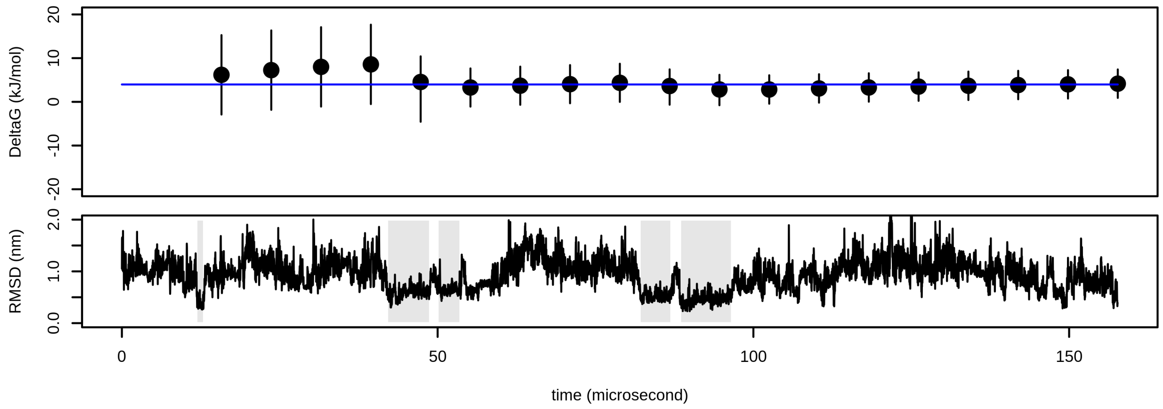


Figure S23: Confidence intervals of folding free energy of Protein B. The value of free energy calculated for the whole simulation is depicted as a blue line.

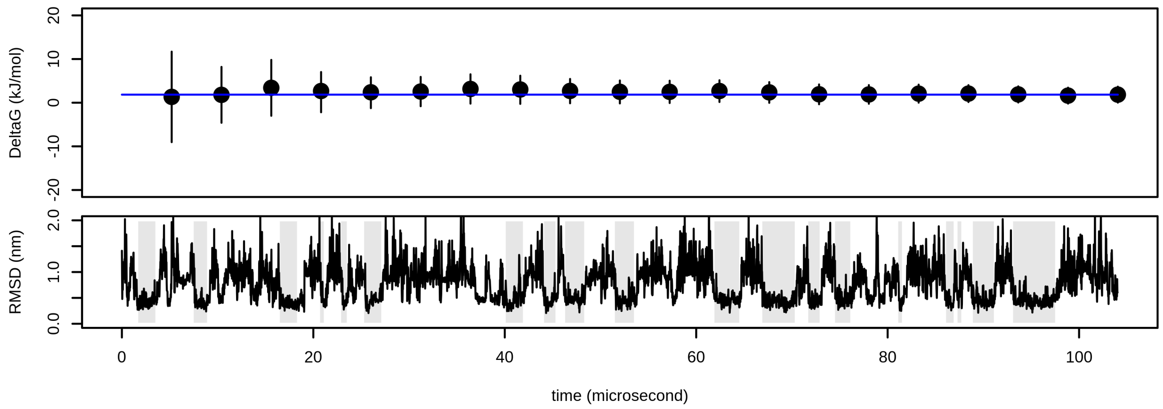

Figure S24: Confidence intervals of folding free energy of Homeodomain (simulation 0). The value of free energy calculated from all simulations is depicted as a blue line.

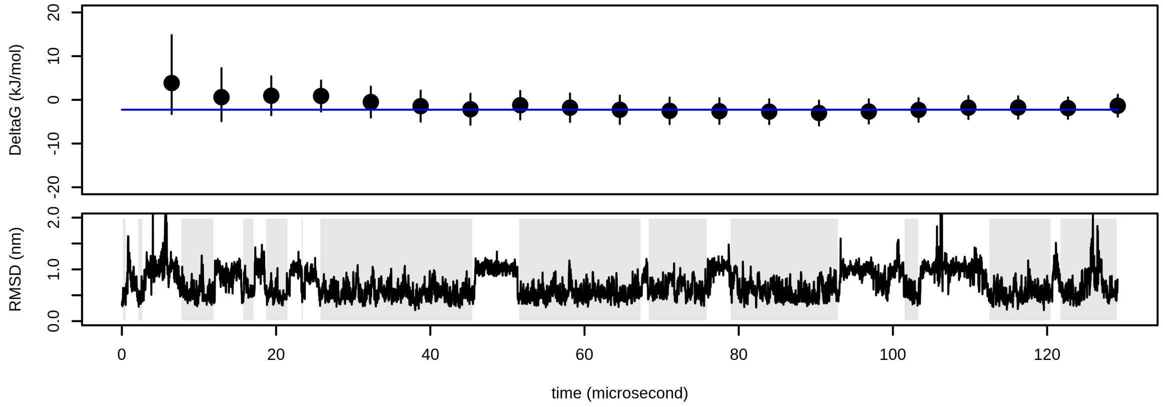


Figure S25: Confidence intervals of folding free energy of Homeodomain (simulation 1). The value of free energy calculated from all simulations is depicted as a blue line.

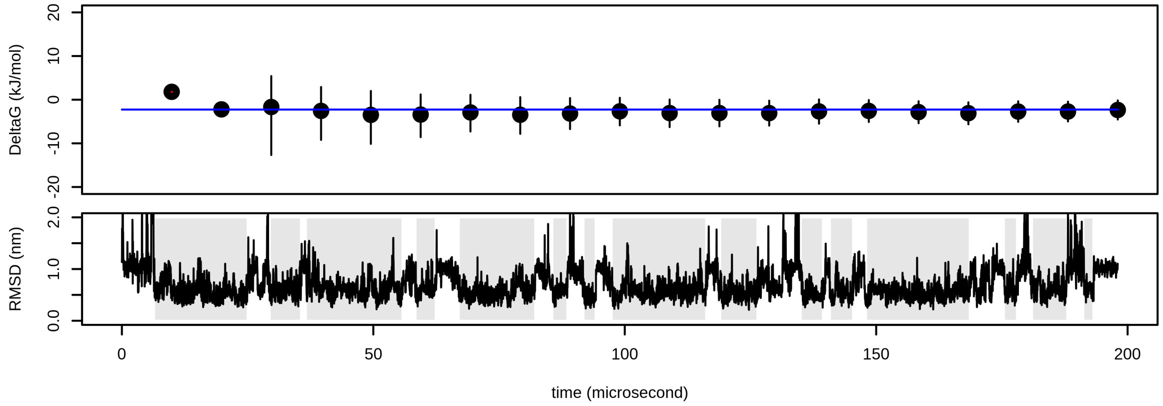

Figure S26: Confidence intervals of folding free energy of Protein G (simulation 0). The value of free energy calculated from all simulations is depicted as a blue line. A confidence interval that does not span this value is depicted in red.

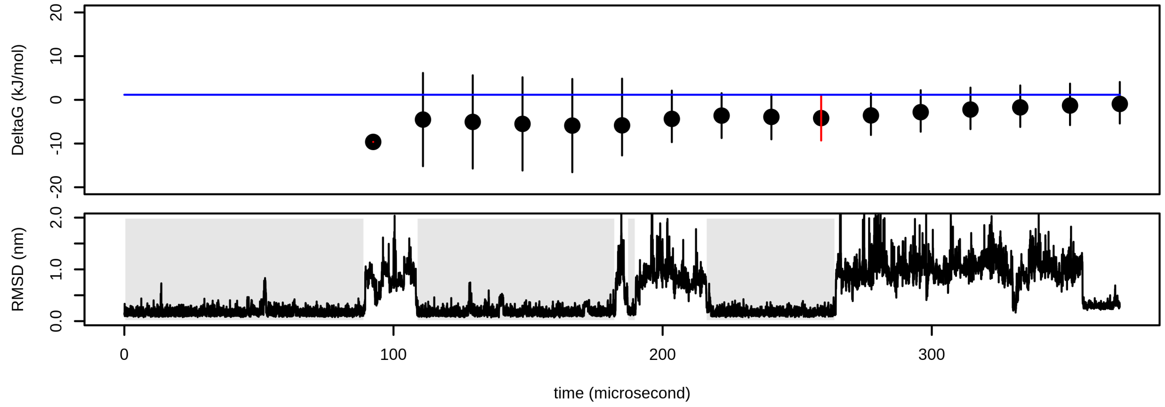


Figure S27: Confidence intervals of folding free energy of Protein G (simulation 1). The value of free energy calculated from all simulations is depicted as a blue line.

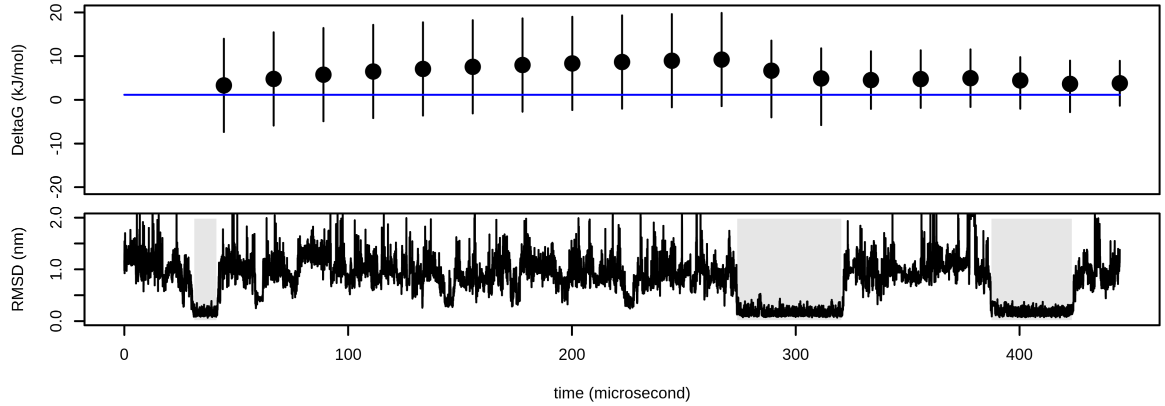

Figure S28: Confidence intervals of folding free energy of Protein G (simulation 2). The value of free energy calculated from all simulations is depicted as a blue line.

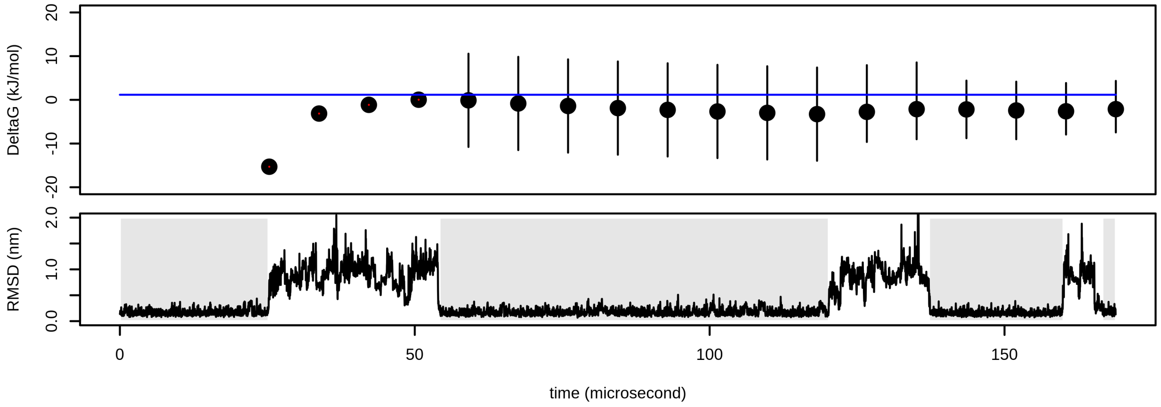


Figure S29: Confidence intervals of folding free energy of Protein G (simulation 3). The value of free energy calculated from all simulations is depicted as a blue line.

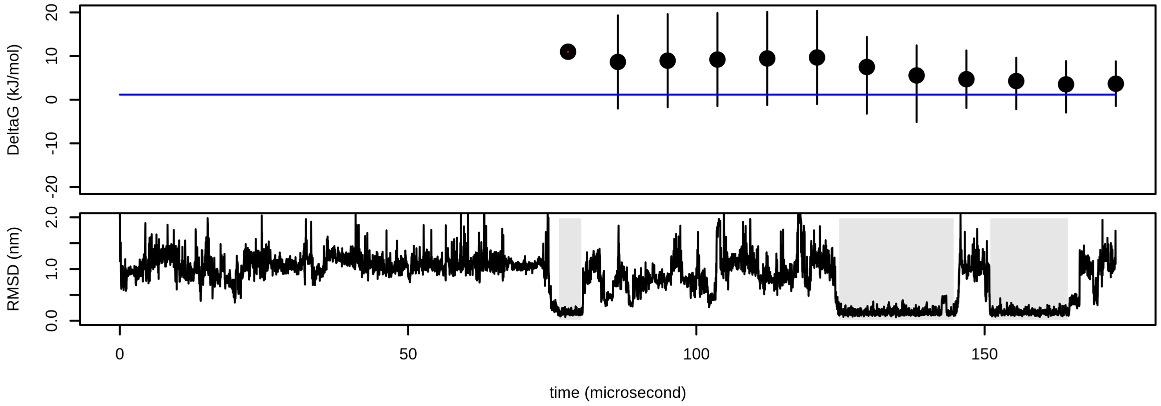

Figure S30: Confidence intervals of folding free energy of $\alpha$3D (simulation 0). The value of free energy calculated from all simulations is depicted as a blue line.

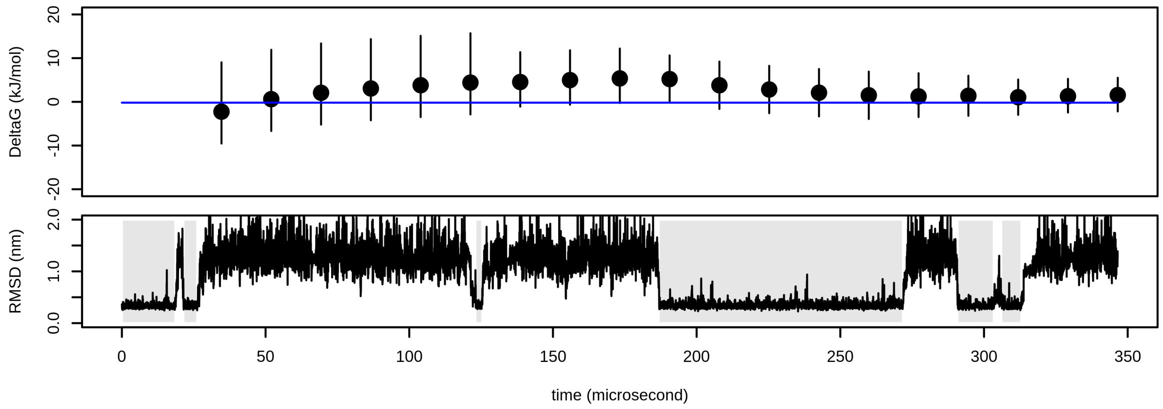


Figure S31: Confidence intervals of folding free energy of $\alpha$3D (simulation 1). The value of free energy calculated from all simulations is depicted as a blue line. Confidence intervals that do not span this value are depicted in red.

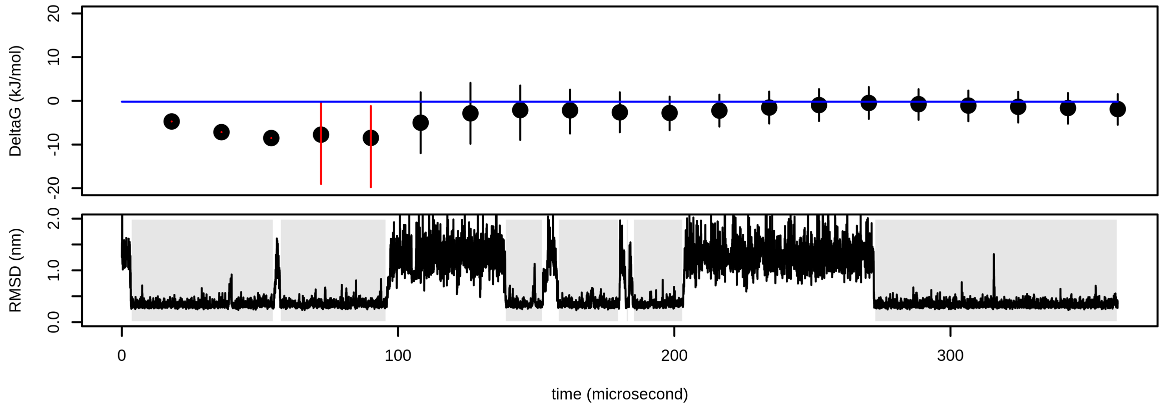

Figure S32: Confidence intervals of folding free energy of $\lambda$-repressor (simulation 0). The value of free energy calculated from all simulations is depicted as a blue line.

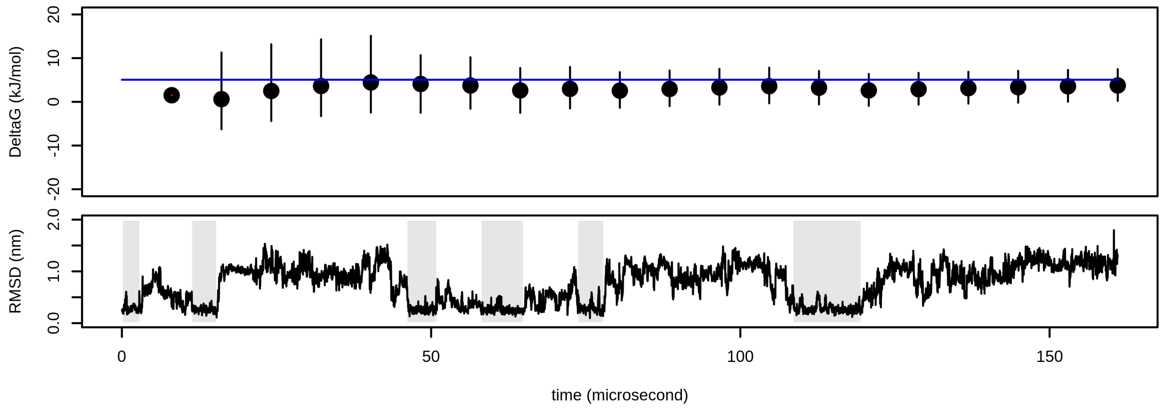


Figure S33: Confidence intervals of folding free energy of $\lambda$-repressor (simulation 1). The value of free energy calculated from all simulations is depicted as a blue line.

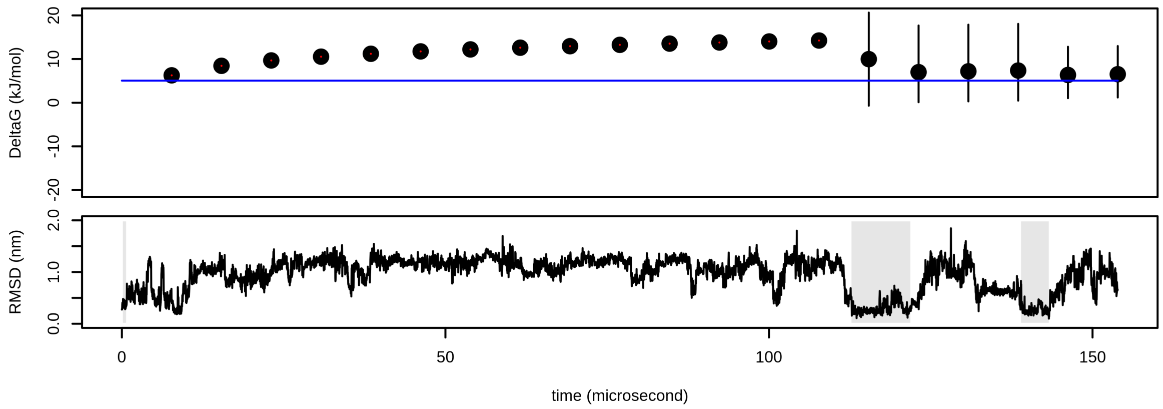

Figure S34: Confidence intervals of folding free energy of $\lambda$-repressor (simulation 2). The value of free energy calculated from all simulations is depicted as a blue line.

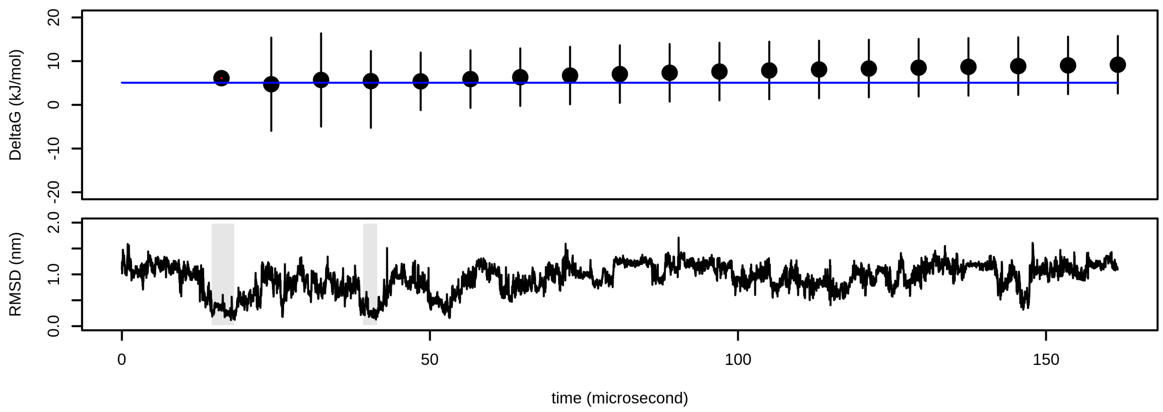


Figure S35: Confidence intervals of folding free energy of $\lambda$-repressor (simulation 3). The value of free energy calculated from all simulations is depicted as a blue line.

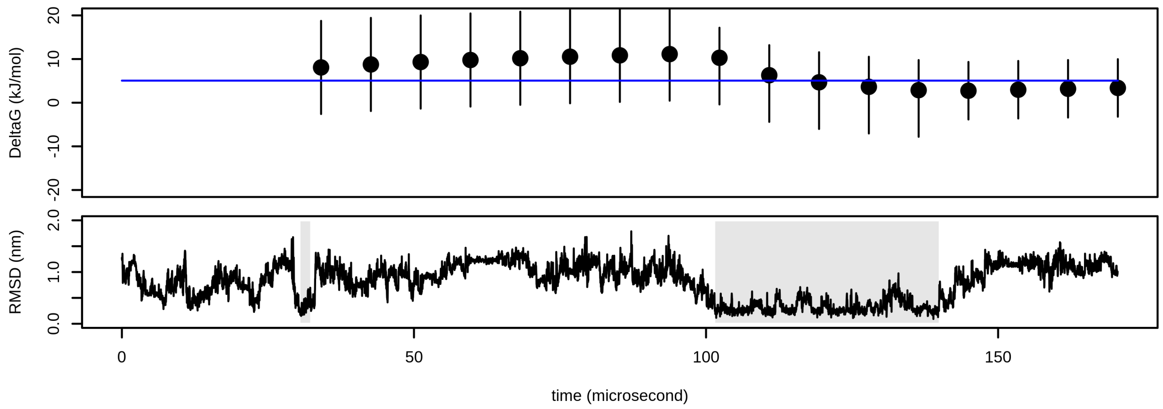

### Calculation of errors in various programming and statistical languages

## R

In R, 95-% CI for $\Delta G$ can be calculated as:

```
nA <- 3       # number of A to B transitions
nB <- 3       # number of B to A transitions
alpha <- 0.05 # value of significance level (0.05 for 95 % CI)
temp <- 300   # temperature in Kelvins
err_top    <-  8.314*temp*log(qf(p=(1-alpha/2), df1=2*nB, df2=2*nA))
err_bottom <- -8.314*temp*log(qf(p=alpha/2, df1=2*nB, df2=2*nA))
print(err_top/1000)     # height of top errorbar in kJ/mol
print(err_bottom/1000)  # height of bottom errorbar in kJ/mol
```

#### Python

In Python (with package `scipy`), 95-% CI for $\Delta G$ can be calculated as:

```
import scipy as sp
from scipy.stats import f
nA = 3         # number of A to B transitions
nB = 3         # number of B to A transitions
alpha = 0.05   # value of significance level (0.05 for 95 % CI)
temp = 300.0   # temperature in Kelvins
err_top    =  8.314*temp*sp.log(f.ppf(q=(1-alpha/2), dfn=2*nB, dfd=2*nA))
err_bottom = -8.314*temp*sp.log(f.ppf(q=alpha/2, dfn=2*nB, dfd=2*nA))
print(err_top/1000.0)     # height of top errorbar in kJ/mol
print(err_bottom/1000.0)  # height of bottom errorbar in kJ/mol
```

#### Wolfram Mathematica

In Wolfram Mathematica, 95-% CI for $\Delta G$ can be calculated as:

```
nA = 3;         (* number of A to B transitions*)

nB = 3;         (*number of B to A transitions*)

alpha = 0.05;   (*value of significance level (0.05 for 95 % CI)*)

temp = 300.0;   (* temperature in Kelvins*)

errTop = 8.314*temp*Log[Quantile[FRatioDistribution[2*nB, 2*nA], 1 - alpha/2]];

errBottom = -8.314*temp*Log[Quantile[FRatioDistribution[2*nB, 2*nA], alpha/2]];

Print[errTop/1000.]     (*height of top errorbar in kJ/mol*)

Print[errBottom/1000.]  (*height of bottom errorbar in kJ/mol*)
```

#### Octave

In Octave, 95-% CI for $\Delta G$ can be calculated as:

```
nA = 3           # number of A to B transitions

nB = 3           # number of B to A transitions

alpha = 0.05     # value of significance level (0.05 for 95 % CI)

temp = 300.0     # temperature in Kelvins

errTop = 8.314*temp*log(finv(1 - alpha/2, 2*nB, 2*nA))

errBottom = -8.314*temp*log(finv(alpha/2, 2*nB, 2*nA))

errTop/1000.     # height of top errorbar in kJ/mol

errBottom/1000. # height of bottom errorbar in kJ/mol
```

#### Julia

In Julia (with package `Distributions`), 95-% CI for $\Delta G$ can be calculated as:

```
using Distributions

nA = 3         # number of A to B transitions

nB = 3         # number of B to A transitions

alpha = 0.05   # value of significance level (0.05 for 95 % CI)

temp = 300.0   # temperature in Kelvins

d = FDist(2*nB,2*nA)

err_top    =  8.314*temp*log(quantile(d, 1-alpha/2.0))

err_bottom = -8.314*temp*log(quantile(d, alpha/2.0))

print(err_top/1000.0)     # height of top errorbar in kJ/mol

print(err_bottom/1000.0)  # height of bottom errorbar in kJ/mol
```
